# Effect of RBD mutation (Y453F) in spike glycoprotein of SARS-CoV-2 on neutralizing antibody affinity

**DOI:** 10.1101/2020.11.27.401893

**Authors:** Takuma Hayashi, Nobuo Yaegashi, Ikuo Konishi

## Abstract

Natural selection “adaptation” in the coronavirus can occur during coronavirus amplification *in vivo* in farmed minks. Natural selection in such viruses is observed by introduction of mutations in SARS-CoV-2 that are not observed during the growth process in humans. Infection with a mutant (Y453F) of SARS-CoV-2 from farmed minks is known to widely spread among humans. We investigated the virological characteristics of this SARS-CoV-2 mutant (Y453F) using three-dimensional protein structural analysis. Our experimental study suggests that virus variants with the Y453F mutation partially escaped detection by four neutralizing monoclonal antibodies. The spread of SARS-CoV-2 variants mediated by millions of infected farmed minks is uncontrolled; consequently, raising a concern that infection of SARS-CoV-2 mutants that cause serious symptoms in humans may spread globally.

## Introduction

Mutations in severe acute respiratory syndrome coronavirus-2 (SARS-CoV-2) can jeopardize the efficacy of potential vaccines and therapeutics against coronavirus disease-2019 (COVID-19). Animals of the Mustelidae family, such as minks and ferrets can be infected with SARS-CoV-2 relatively easily compared with other mammals (1). However, the reason why SARS-CoV-2 is extremely contagious to these animals remains to be elucidated. Nonetheless, it is clear that when several farmed minks kept in a high-density environment are infected with SARS-CoV-2, the virus proliferates in large numbers. Consequently, humans and minks may be at high risk of SARS-CoV- 2 infection.

Natural selection “adaptation” in the coronavirus can occur during coronavirus amplification *in vivo* in farmed minks (2). Natural selection in such viruses is observed by introduction of mutations in SARS-CoV-2 that are not observed during the growth process in humans (2,3). Infection with a mutant (Y453F) of SARS-CoV-2 from farmed minks is known to widely spread among humans (4). We investigated the virological characteristics of this SARS-CoV-2 mutant (Y453F) using threedimensional protein structural analysis.

## Methods

SARS-CoV-2 mutant has an amino acid mutation Y453F in the sequence encoding spike glycoprotein (4). This SARS-CoV-2 mutant has been detected in approximately 300 viral sequences isolated from the European and Dutch population as well as in minks.

Data on the three-dimensional structure of the receptor binding domain (RBD) of the spike glycoprotein of SARS-CoV-2 (PDB ENTITY SEQ 6VW1_1) was used in these studies (5). Data (PDB: 6XC2, 6XC4, 7JMP, 7JMO, 6XKQ, and 6XKP) on the three-dimensional structure of six neutralizing antibodies (CC12.1, CC12.3, COVA2-39, COVA2-04, CV07-250, and CV07-270) that bind to the spike glycoprotein of SARS-CoV-2 was used in these studies (6).

Using the Spanner program, we predicted the three-dimensional structure of the SARS-CoV-2 spike glycoprotein Y453F mutant based on PDB (ENTITY SEQ 6VW1_1). We investigated the binding of the spike glycoprotein Y453F mutant of SARS-CoV-2 to human angiotensin-converting enzyme 2 (ACE2) and determined the affinity of the spike glycoprotein Y453F mutant of SARS-CoV-2 to six neutralizing monoclonal antibodies using the MOE program (three-dimensional protein structure modeling, protein docking analysis: MOLSIS Inc., Tokyo, Japan) and Cn3D macromolecular structure viewer.

## Results

The Y453F mutation did not affect the three-dimensional structure of conventional SARS-CoV-2 spike glycoproteins. From the present study, it was clarified that the binding property of the spike glycoprotein Y453F mutant to human ACE2 was slightly weaker than that of the conventional SARS- CoV-2 spike glycoprotein (Figure 1A). This was due to the replacement of Tyr at position 453 by Phe, which was unable to form a hydrogen bond with His at position 34 in human ACE2.

**Figure 1:**
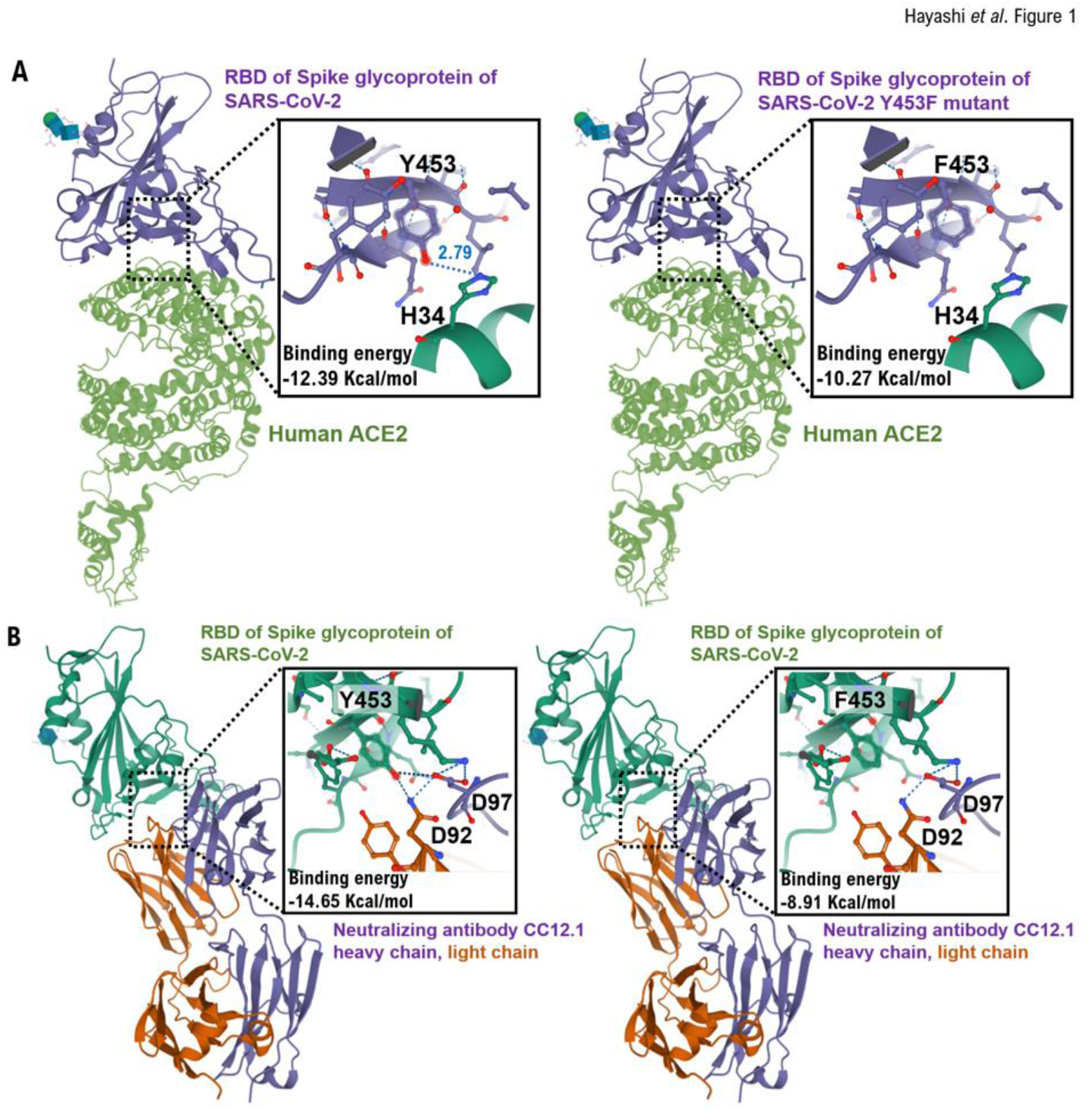
Interactions between the RBD and RBD Y453F mutant in the spike glycoprotein of SARS-CoV-2 and heavy chain of neutralizing monoclonal antibody (CC12.2). **(A)** The interaction between Angiotensin-converting enzyme 2 (ACE2) (green) and residues of the conventional receptor binding domain (RBD) or RBD Y453F mutant (purple) is shown using the three-dimensional structure model. It is speculated that the Y453 amino acid residue of the conventional RBD is hydrogen-bonded to the H34 amino acid residue of human ACE2. However, the binding between the F453 amino acid residue of the RBD mutant and the H34 amino acid residue of human ACE2 is presumed to be slightly weak. From these results, the affinity between the spike glycoprotein of the RBD Y453F mutant and human ACE2 is presumed to be slightly weak compared with the conventional RBD. **(B)** The interaction between the heavy chain (purple) and light chain (brown) of neutralizing monoclonal antibody CC12.1 and residues of the conventional RBD or RBD Y453F mutant (green) is shown using the three-dimensional structure model. It is speculated that the Y453 amino acid residue of the conventional RBD is hydrogen-bonded to the D92 amino acid residue of the light chain and D97 amino acid residue of the heavy chain of neutralizing monoclonal antibody CC12.1. However, the binding between the F453 amino acid residue of the RBD mutant and the D92 and D97 amino acid residues of neutralizing monoclonal antibody CC12.1 is presumed to be weak. From these results, the affinity between the spike glycoprotein of the RBD Y453F mutant and neutralizing monoclonal antibody CC12.1 is presumed to be low compared with the conventional RBD. The three-dimensional structure models are shown by Cn3D macromolecular structure viewer.

The present study revealed that the affinity between the spike glycoprotein Y453F mutant and four of the six monoclonal antibodies (CC12.1, CC12.3, COVA2-39, COVA2-04, CV07-250, CV07-270) examined was clearly weak compared with the conventional SARS-CoV-2 spike glycoprotein (Figure 1B, Table 1).

**Table 1.** Binding energy between the RBD of the spike glycoprotein of SARS-CoV-2 and neutralizing monoclonal antibodies. The binding of the spike glycoprotein Y453F mutant of SARS-CoV-2 to human ACE2 and the affinity of the spike glycoprotein Y453F mutant of SARS-CoV-2 to six neutralizing monoclonal antibodies were investigated using the MOE program (three-dimensional protein structure modeling, protein docking analysis: MOLSIS Inc., Tokyo, Japan) and Cn3D macromolecular structure viewer. Binding energy calculated by the MOE program is shown in the table.

| Binding energy between RBD and human ACE2 |  | Effect of mutation on antibody |
| --- | --- | --- |
| RBD | RBR (Y453F) |  |
| -12.39 Kcal/mol | -10.27 Kcal/mol | Slightly lower |

| Neutralizing antibody | Binding energy between RBD and antibodies |  | Effect of mutation on antibody |
| --- | --- | --- | --- |
|  | RBD | RBD (Y453F) |  |
| CC12.1 | -14.65 Kcal/mol | -8.91 Kcal/mol | Low affinity |
| CC12.3 | -5.85 Kcal/mol | -1.36 Kcal/mol | Low affinity |
| COVA2-39 | not calculated | not calculated | NA |
| COVA2-04 | -12.4 Kcal/mol | -3.42 Kcal/mol | Low affinity |
| CV07-250 | -1.09 Kcal/mol | -0.23 Kcal/mol | Low affinity |
| CV07-270 | not calculated | not calculated | NA |

| Binding energy between RBD and human ACE2 |  |  | Effect of mutation on antibody |
| --- | --- | --- | --- |
| RBD | RBR (Y453F) |  |  |
| -12.39 Kcal/mol | -10.27 Kcal/mol |  | Slightly lower |
| Neutralizing antibody | Binding energy between RBD and antibodies |  | Effect of mutation on antibody |
|  | RBD | RBD (Y453F) |  |
| CC12.1 | -14.65 Kcal/mol | -8.91 Kcal/mol | Low affinity |
| CC12.3 | -5.85 Kcal/mol | -1.36 Kcal/mol | Low affinity |
| COVA2-39 | not calculated | not calculated | NA |
| COVA2-04 | -12.4 Kcal/mol | -3.42 Kcal/mol | Low affinity |
| CV07-250 | -1.09 Kcal/mol | -0.23 Kcal/mol | Low affinity |
| CV07-270 | not calculated | not calculated | NA |

It is considered that the affinity between the appropriate amino acid residues in the variable region of the antibody and the spike glycoprotein Y453F mutant was diminished owing to weak recognition of the monoclonal antibody to spike glycoproteins.

## Discussion

To the best of our knowledge, data on all SARS-CoV-2 mutants have not been published till date. Therefore, it is unclear whether SARS-CoV-2 mutants detected in people working on mink farms are actually derived from the farmed minks. However, in the present study the subspecies of SARS-CoV- 2 derived from farmed minks has been observed in the group of infected people, and the virus mutants that were inherited by infected individuals.

Mutations in SARS-CoV-2 that lead to generation of SARS-CoV-2 subspecies, have made humans and animals susceptible to infection through easy propagation in the host, thereby making it difficult to identify the effects of therapeutic agents or vaccines for COVID-19. Moreover, the spread of SARS-CoV-2 variants mediated by millions of infected farmed mink is uncontrolled; consequently, raising a concern that infection of SARS-CoV-2 mutants that cause serious symptoms in humans may spread globally.

## Data Sharing

Data are available on various websites and have also been made publicly available (more information can be found in the first paragraph of the Results section).

## Acknowledgments

We thank Professor Richard A. Young (Whitehead Institute for Biomedical Research, Massachusetts Institute of Technology, Cambridge, MA) for his research assistance. This study was supported in part by grants from the Japan Ministry of Education, Culture, Science and Technology (No. 24592510, No. 15K1079, and No. 19K09840); Foundation of Osaka Cancer Research; Ichiro Kanehara Foundation for the Promotion of Medical Sciences and Medical Care; Foundation for Promotion of Cancer Research; Kanzawa Medical Research Foundation; Shinshu Medical Foundation; and Takeda Foundation for Medical Science.

## Author Contributions

T.H. performed most of the experiments and coordinated the project. T.H. and N.Y. conceived the study and wrote the manuscript. N.Y. and I.K. provided with information on clinical medicine and oversaw the entire study.

## Transparency Document

The transparency document associated with this article can be found in the online version at http://.

